# Characterization of focused ultrasound blood-brain barrier disruption effect on inflammation as a function of treatment parameters

**DOI:** 10.1101/2024.07.10.602776

**Authors:** Cleide Angolano, Emily Hansen, Hala Ajjawi, Paige Nowlin, Yongzhi Zhang, Natalie Thunemann, Christiane Ferran, Nick Todd

## Abstract

The technology of focused ultrasound-mediated disruption of the blood-brain barrier (FUS-BBB opening) has now been used in over 20 Phase 1 clinical trials to validate the safety and feasibility of BBB opening for drug delivery in patients with brain tumors and neurodegenerative diseases. The primary treatment parameters, FUS intensity and microbubble dose, are chosen to balance sufficient BBB disruption to achieve drug delivery against potential acute vessel damage leading to microhemorrhage. This can largely be achieved based on both empirical results from animal studies and by monitoring the microbubble cavitation signal in real time during the treatment. However, other safety considerations due to second order effects caused by BBB disruption, such as inflammation and alteration of neurovascular function, are not as easily measurable, may take longer to manifest and are only beginning to be understood. This study builds on previous work that has investigated the inflammatory response following FUS-BBB opening. In this study, we characterize the effect of FUS intensity and microbubble dose on the extent of BBB disruption, observed level of microhemorrhage, and degree of inflammatory response at three acute post-treatment time points in the wild-type mouse brain. Additionally, we evaluate differences related to biological sex, presence and degree of the anti-inflammatory response that develops to restore homeostasis in the brain environment, and the impact of multiple FUS-BBB opening treatments on this inflammatory response.

## Introduction

Focused ultrasound (FUS)-mediated disruption of the blood-brain barrier (BBB) has been safely implemented in multiple Phase 1 clinical trials^1–3^. This platform technology for transiently increasing the permeability of the BBB to improve drug penetrance to the brain has the potential to be used in a wide range of neurological disorders^4–7^. Current clinical treatments are being done safely, as defined by a successful extravasation of MRI contrast agent, absence of tissue damage on post-treatment MRI and no reported adverse events. Balancing FUS parameters to achieve sufficient BBB disruption without causing acute damage such as microhemorrhage can largely be done through monitoring the cavitation signal during treatment. However, second order effects due to BBB disruption, including induction of a neuroinflammatory response, cannot be easily monitored, may take longer to manifest and are not as well understood^8,9^. Expanding treatments to new indications in other neurological disorders will likely require more aggressive BBB opening to deliver larger therapeutics such as biologics and gene therapies, BBB disruption over larger brain volumes, and repeated treatments. For certain patient populations these treatments will need to occur in the context of diseased brains with pre-existing neuroinflammation. To safely advance the field into new indications, the impact of FUS-BBB opening on the brain inflammatory response needs to be better characterized and understood.

In 2017, Kovacs et al. demonstrated that FUS-BBB opening using a high dose of microbubbles in the rat brain led to a transient upregulation of several inflammatory markers, similar to what was reported as a sterile inflammatory response in mild traumatic brain injuries^10^. Several studies since in healthy wild type mice further characterized the inflammatory response induced by FUS-BBB opening in terms of changes in the brain transcriptome (RNA), proteome, and levels of trophic factors, time course of this response, and its dependence on FUS treatment parameters^10–15^. Most studies identified increased astrocytosis and microgliosis, as well as upregulation of cytokines, chemokines and adhesion molecules in the first 24-hours post-treatment, as hallmarks of the inflammatory response. **Figure 1** summarizes the findings from 6 papers linking FUS parameters to the intensity of the inflammatory response. Collectively, these findings suggest that a significant inflammatory response is evident at higher microbubble dose or stronger FUS intensity. However, a window of treatment parameter combinations in which BBB opening can be achieved without inducing a significant inflammatory response does exist.

**Figure 1.**
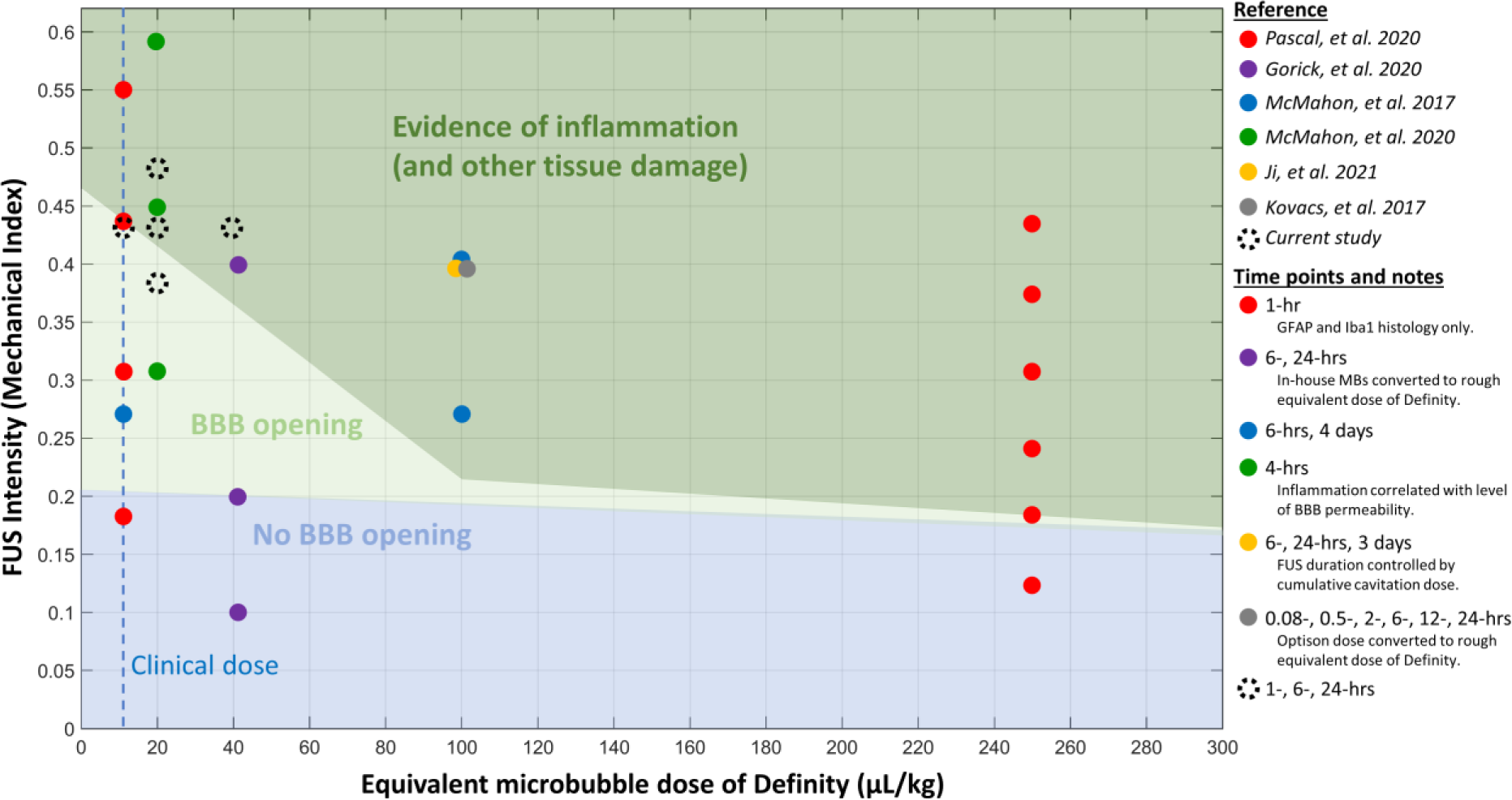
Literature reports of BBB opening and inflammation as a function of FUS intensity and microbubble dose treatment parameters. Colored circles represent experimental groups at a particular FUS intensity and microbubble dose combination for each given publication. Background shaded regions indicate whether the publication reported no BBB opening, BBB opening only, or BBB opening plus evidence of an inflammatory response for the treatment parameters used. Colored circles on a boundary indicate results were equivocal for this parameter combination. Dashed black circles represent the treatment parameters considered in this study. The vertical dashed blue line indicates the currently approved clinical dose level of 10 µL/kg of Definity.

The goal of this study is to build upon the existing results by 1) characterizing the inflammatory response for a range of FUS intensity and microbubble dose combinations that are near the boundary of inflammation onset; 2) gauging whether there are differences in the inflammatory response related to sex; 3) identifying the protective components of the response to injury whose function is to bring the brain environment back to homeostasis; and 4) evaluating the influence of multiple FUS-BBB opening treatments on the severity of the inflammatory response.

## Methods

### Experimental Design

All experiments were carried out under approval from the Brigham and Women’s Hospital Institutional Animal Care and Use Committee. 130 CD-1 mice (25 – 30g, Charles River Laboratories, Wilmington, MA) were used in 20 experimental groups to characterize the response to FUS-BBB opening treatment as a function of FUS intensity, microbubble dose, and time from treatment (Table 1). In the first set of studies evaluating incremental FUS intensities or microbubble doses, mice received one treatment of FUS-BBB opening. FUS intensities of 0.32, 0.36 and 0.40 MPa were used with a single microbubble dose of 20 µL/kg.

**Table 1.** Experimental groups used in this study. For each group, the FUS intensity, microbubble dose, post- treatment sacrifice time, and number of mice are noted.

| Experimental Groups |  | Time to sacrifice |  |  |  |
| --- | --- | --- | --- | --- | --- |
|  |  | 1 hr | 6 hr | 24 hr | No FUS |
| **FUS intensity**<br>• 20 $\mu\text{L/kg}$ Definity bubble dose<br>• 1 target location | 0.32 MPa | N=8 | N=8 | N=8 | N=6 |
|  | 0.36 MPa * | N=6 | N=6 | N=6 | N=6 |
|  | 0.40 MPa | N=6 | N=6 | N=6 | -- |
| **Microbubble dose**<br>• 0.36 MPa<br>• 1 target location | 10 $\mu\text{L/kg}$ | N=6 | N=6 | N=6 | N=6 |
| | 20 $\mu\text{L/kg}$ * | N=6 | N=6 | N=6 | N=6 |
| | 40 $\mu\text{L/kg}$ | N=6 | N=6 | N=6 | N=6 |
| **5x Multiple Treatments**<br>• 20 $\mu\text{L/kg}$ Definity bubble dose<br>• 1 target location | 0.32 MPa | -- | -- | N=8 | -- |

\* 0.36 MPa, 20 $\mu\text{L/kg}$ microbubble dose mouse groups were performed once and used in both studies

Microbubble doses of 10, 20 and 40 µL/kg were used with a single FUS intensity of 0.36 MPa. The mouse group treated at 0.36 MPa FUS intensity and 20 µL/kg microbubble dose was done once and used in both sets of analyses. In each treatment group, mice were sacrificed at 1-, 6-, and 24-hours post-treatment. An equal number of male and female mice were used in all experimental groups. In the second set of experiments aimed at investigating the effects of multiple FUS-BBB opening treatments, a group of 8 mice (all males) received five sequential weekly treatments of FUS-BBB opening performed at 0.32 MPa FUS intensity and 20 µL/kg microbubble dose. Mice were sacrificed 24 hours after the final treatment.

### FUS Blood-Brain Barrier Opening

FUS-BBB opening treatments were performed with an in-house built FUS system consisting of a 690 kHz single element geometrically focused transducer, passive cavitation detector and manual three-axis positioning system. Mice were anesthetized with 100 mg/kg ketamine and 10 mg/kg xylazine. Their heads were shaved and any remaining hair was removed with depilatory cream. An IV catheter was placed in the tail vein to administer the microbubbles. Mice were placed in a positioning jig with fixation points at ear bars and teeth, allowing for reproducible positioning relative to the transducer. The focus of the transducer was set at 2.5 mm lateral and 4.0 mm anterior to the interaural line, targeting the right striatum. Microbubble injections at the specified dose were given by bolus tail vein injection immediately prior to the onset of FUS sonications. Sonications at a single target were performed at the specified FUS intensity in 10 ms bursts and 1 Hz repetition frequency for 2 minutes. The dye Trypan blue (4% concentration, 2.5 ml/kg) was administered by tail vein injection immediately after FUS sonication. Mice were recovered on a warm pad and re-housed until sacrifice.

### Mouse sacrifice and tissue harvest

Following isoflurane anesthesia, mice were sacrificed by cervical dislocation, and their fresh brains recovered and placed in a mouse brain matrix (Electron Microscopy Sciences, Road Hatfield, PA). Using the visible trypan blue dye as a guide, three cuts were made to acquire two 2mm slabs of tissue (see **Figure S1**). The anterior slab was split into ipsilateral (FUS-targeted) and contralateral (non-targeted) hemispheres and individually snap frozen in liquid nitrogen. These tissue samples were stored at -80°C for subsequent RNA and protein isolation. The posterior slab was processed for immunohistochemistry (IHC) and immunofluorescence (IF) analysis. All reagents used in this study are listed in Supplementary Materials.

### Staining and Immunohistochemistry (IHC)

Brain sections were processed for staining, IHC and IF as described in Guedes et al. 2014^16^. In brief, 2000 µM-coronal tissue slabs were zinc-fixed (BD Pharmigen, San Diego, CA, USA) for 48h at room temperature, dehydrated in a tissue processor and embedded in paraffin, before being sectioned into 6 µM thickness.

The extent of the BBB-opening following FUS treatment was estimated by measuring the Trypan blue extravasation, using trypan blue autofluorescence properties. Brain sections were scanned using Odyssey CLx (Li-Cor biosciences, Lincoln, NE). Wavelengths were 685 nm for excitation and 710-730 nm for detection. Autofluorescence signals were quantified (area, pixels) using Image StudioTM Software version 5.2.5 (Li-Cor biosciences, Lincoln, NE).

For histological evaluation and quantification of hemorrhage, sections were stained with Wright-Giemsa stain modified kit (Sigma-Aldrich Saint Louis, MO). Sections were scanned using Mica imaging system at 10x magnification (Leica Microsystems Inc., Deerfield, IL) and images were quantified (area, pixels) using ImageJ version 1.53 (NIH, Bethesda, MD).

For IHC, sections were de-paraffinized and rehydrated. Antigen retrieval was performed using citrate buffer 10mM pH 6.0 for 7 minutes at 95-100°C. Sections were then incubated with horse serum (7% in PBS) for 20 minutes prior to overnight incubation at 4°C with goat anti-intracellular adhesion molecule-1 (ICAM1) (R&D Systems, Inc, Minneapolis, MN, USA), rabbit anti-glial fibrillary acidic protein (GFAP, Abcam, Waltham, MA) and rabbit anti-TMEM-119 (Abcam). Sections were then treated with H_2_O_2_ 1:100 in PBS for 10 min, incubated with the appropriate biotinylated secondary IgG antibodies (Vector Laboratories, Burlingame, CA), followed by ABC (avidin-biotin complex) reagent (Vector Laboratories), then detected by ImmPACT 3,3’-diaminobenzidine tetrahydrochloride (DAB) peroxidase substrate (Vector Laboratories). Negative controls using only secondary antibodies confirm the absence of non-specific immunostaining.

### mRNA isolation and cPCR

mRNA was isolated from tissue lysates using RNA/protein purification kit (Norgen Biotek Corp., Thorold, ON, Canada) then cDNA was synthesized using iScript cDNA synthesis kit (Bio-Rad, Hercules, CA, USA). qPCR was performed using iTaq Fast SYBR Green Supermix with ROX (Bio-Rad, Hercules, CA) and gene-specific primers (Integrated DNA Technologies, Coralville, IA and Sigma-Aldrich, St. Louis, MO) (Suppl. Table I), using ABI 7500 Fast Real-Time PCR System (Applied Biosystems, Inc., Foster City, CA). Comparative threshold cycle (Ct) method was used to perform relative quantification of qPCR results. mRNA expression of target genes *Gfap*, *Iba-1*, *Ikba*, *Nlrp3*, *Il1b*, *Il6*, *Il10*, *Ccl2*, *Icam*, *Vcam*, *Mmp9*, *iNos*, *Sele*, *eNos*, *Vegf*, and *A20*/*Tnfaip3* were normalized to that of the housekeeping genes 28S ribosomal RNA, valosin-containing protein (VCP), and TATA binding protein (TBP) using BestKeeper method^17^. Data are expressed as fold change of control mice.

### Western blot analysis

Tissue lysates (30-40ug protein) were separated under reducing conditions by sodium dodecyl sulphate-polyacrylamide gel electrophoresis (SDS-PAGE) (4-20% Criterion TGX Stain-free gels, Bio-Rad Laboratories, Hercules, CA, USA) and transferred to Polyvinylidene fluoride (PVDF) membranes (Trans-Blot Turbo RTA Midi 0.45 µm LF PVDF Transfer Kit, Bio-Rad Laboratories, Hercules, CA, USA) by semi-dry electroblotting (Trans-Blot turbo transfer system, Bio-Rad Laboratories, Hercules, CA, USA). Membranes were probed with rabbit anti-GFAP (Cell Signaling, Danvers, MA, USA), mouse anti-glyceraldehyde 3-phosphate dehydrogenase (GAPDH) (Santa Cruz Biotechnology, Inc, Santa Cruz, CA, USA), and goat anti-intracellular adhesion molecule-1 (ICAM) (R&D Systems, Inc., Minneapolis, MN, USA). Appropriate infrared secondary IgG antibodies IRDye® 680RD and/or IRDye® 800CW (Li-Cor biosciences, Lincoln, NE) were used. Membranes were scanned using Odyssey CLx (Li-Cor biosciences, Lincoln, NE). Densitometric analysis was done using Image StudioTM Software version 5.2.5 (Li-Cor biosciences, Lincoln, NE).

### Statistical analysis

All statistical analyses were performed using GraphPad Prism 10.0.1 software. Primary analysis of all outcome measures for the FUS intensity and microbubble dose studies was performed in the framework of a two-way ANOVA. Separate analyses were performed for the FUS Intensity and Microbubble Dose groups, with main factors for FUS intensity and time, and microbubble dose and time. An F-test was performed to determine whether there were significant effects over the main factors of the ANOVA. For data with significant main effects, Tukey’s multiple comparisons test was performed to test for significant differences between groups. Significance for both main effects and group differences is reported at p < 0.05. Analysis for differences due to sex within one set of FUS parameters were also organized in a two-way ANOVA, with factors for sex and time. Analysis of correlations between BBB opening or hemorrhage and inflammatory markers was performed by calculating the Pearson correlation coefficient. Analysis of the multiple treatments study was performed by comparing the 5x-treated mice (males, 0.32 MPa, 20 µL/kg, 24 hrs sacrifice) against male mice from the 1x-treated group treated with the same FUS parameters (males, 0.32 MPa, 20 µL/kg, 24 hrs sacrifice) using an unpaired, two-tailed t-test.

## Results

### BBB opening as a function of FUS intensity and microbubble dose

Evidence of BBB opening was seen in every mouse that underwent FUS treatment. As expected, a larger area of BBB opening is seen with both greater FUS intensity and higher microbubble dose at all time points. Both the 0.40 MPa and 40 µL/kg groups show significantly greater BBB opening than the 0.32 MPa and 10 µL/kg groups, respectively, and at all time points. The area of trypan blue also increases with time to sacrifice following treatment, likely because of diffusion of the dye through the brain parenchyma over time. Representative images of trypan blue fluorescence signal and the user-defined BBB opening region of interest (ROI, blue outline) are shown for each parameter set and post-treatment sacrifice time point in the FUS Intensity (**Figure 2A**) and Microbubble Dose (**Figure 2B**) cohorts. Data from all groups are quantified as ROI area as a percent of total hemisphere area in the bar plots below the images. Two-way ANOVA analysis indicates significant main effects of both FUS intensity and time for the FUS Intensity groups and significant main effects of both microbubble dose and time for the Microbubble Dose groups. Significant differences between individual groups based on post-hoc analysis are indicated in the graphs.

**Figure 2.**
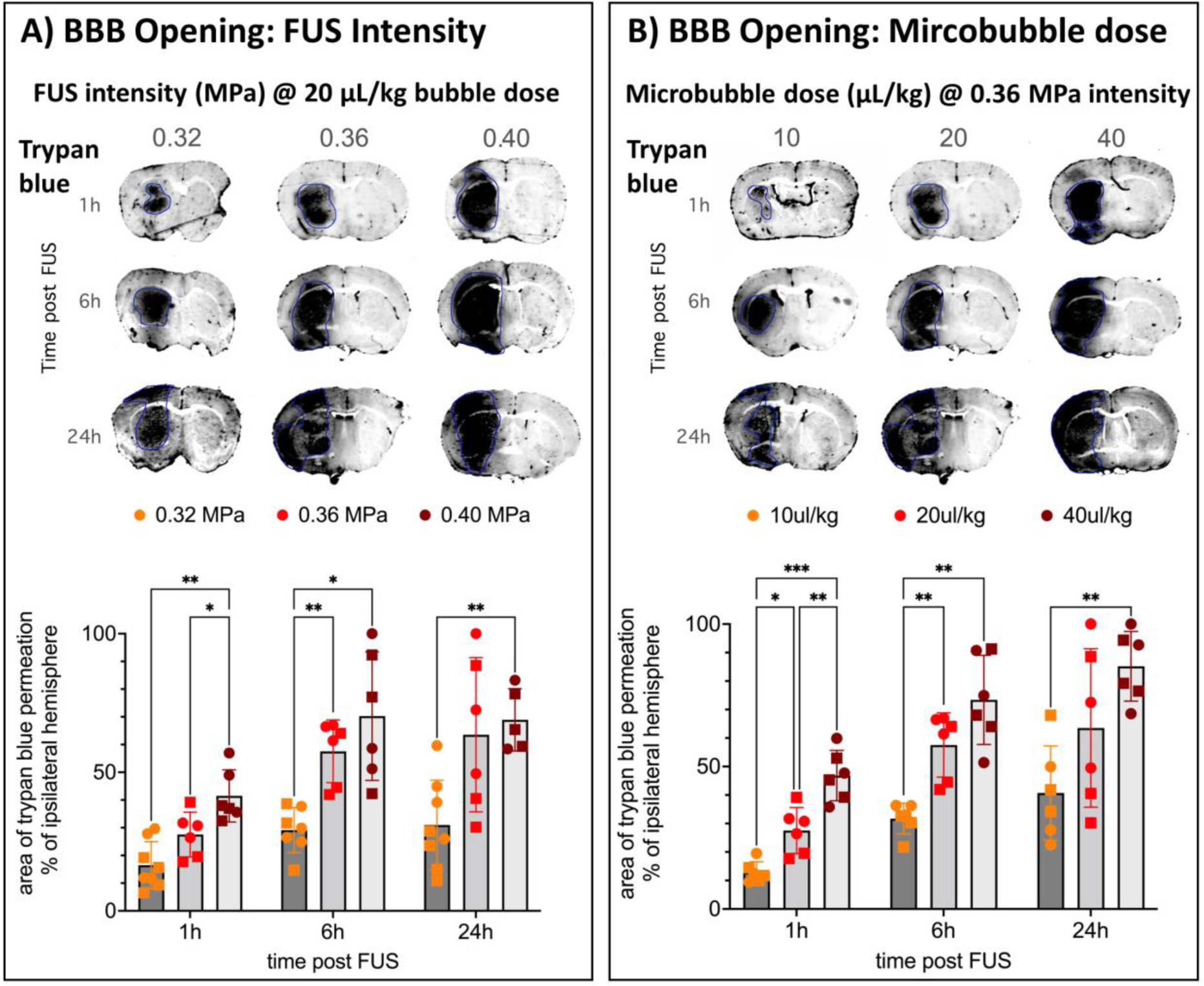
Trypan Blue fluorescence imaging for quantification of BBB opening area. A) Example images of trypan blue fluorescence imaging following FUS-BBB opening for each FUS intensity and time point are shown on top. Blue outlines delineate ROIs drawn to quantify area of BBB opening. Area of BBB opening as a percentage of total hemisphere area are shown in the bar graphs. Significant differences between groups are based on Two-way ANOVA analysis with Tukey’s multiple comparisons test (* = p<0.05, ** = p<0.01, *** = p<0.001). B) Similar example images and bar graphs quantifying BBB opening for the microbubble dose groups. Male mice are plotted as squares, female mice as circles

### Evidence of vessel and tissue damage

FUS BBB opening at lowest intensity and microbubble dose parameters did not cause any microhemorrhage in most mice treated. Only one mouse out of 24 from the 0.32 MPa groups showed any evidence of microhemorrhage and only five out of 24 mice showed signs of hemorrhage at the lowest microbubble dose of 10 µL/kg. However, there was considerable variability in the 0.36 and 0.40 MPa FUS intensity and 20 and 40 µL/kg microbubble dose treatment groups, ranging from no evidence of hemorrhage up to widespread hemorrhage. Results evaluating the presence of microhemorrhage assessed by Wright-Giemsa staining are shown in **Figure 3**. Cell nuclei are stained in purple, cytoplasm in light pink and red blood cells in dark pink to red. Representative images of the FUS-targeted right striatum from the FUS Intensity groups are shown in **Figure 3A**. Bar plots of the hemorrhage area after log transformation and consolidation over time points are shown in **Figure 3B**. Two-way ANOVA analysis of the hemorrhage area for the FUS Intensity groups indicates a highly significant main effect of FUS intensity (p < 0.0001). Post-hoc analysis for significant differences between individual groups are shown in the plots. Similar results for the microbubble dose groups are shown in **Figures 3C – D**. Two-way ANOVA analysis indicates a significant effect of microbubble dose (p = 0.012).

**Figure 3.**
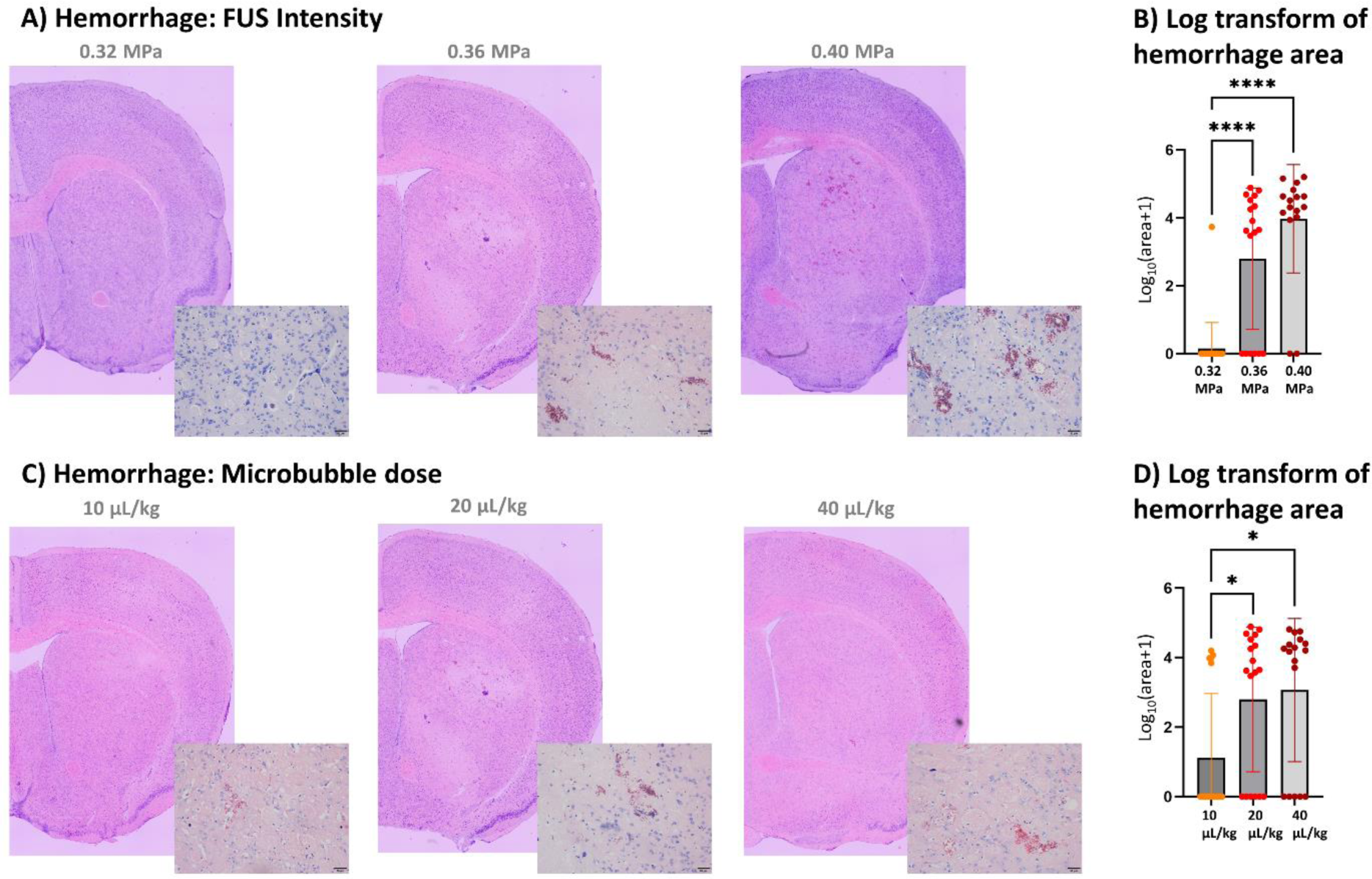
Giemsa staining for quantification of microhemorrhage. A) Representative images of giemsa-stained brain sections showing the FUS-targeted right striatum for mice treated at 0.32, 0.36 and 0.40 MPa FUS intensity. Insets show areas of hemorrhage acquired at higher resolution. B) log10 transform of microhemorrhage area, collapsed over time points. C – D) Similar representative images and bar graphs quantifying microhemorrhage area for the microbubble dose groups. (* = p<0.05, **** = p<0.0001).

In addition to microhemorrhage, we also looked for signs of neuronal cell degeneration based on fluoro-jade staining. Brain sections from two mice (one male and one female) of each group were stained and imaged. All sections were completely negative with no evidence for fluorescent signal (data not shown). This indicates that any damage done to vascular structures was not significant enough to cause neuronal cell death at the 1-, 6- and 24-hr early time points used in this study.

### Pro-inflammatory markers

The heat maps presented in **Figure 4** summarize the mRNA levels measured by qPCR of multiple pro-inflammatory markers in the different treatment groups. For the FUS intensity groups (Figure 4A), only GFAP, a marker of astrocytic activation^18^, demonstrated significant time and intensity-dependent upregulation. GFAP upregulation started 1h after treatment and continued to increase with time. The highest expression levels of GFAP were observed at 24h- post FUS at the highest 0.4 MPa intensity (5.3-fold increase over baseline control). Iba-1, a marker of microglia activation^19^, demonstrated similar gene expression kinetics to GFAP, i.e. peak levels at 24h, however at much lower fold change (∼1.2-fold compared to control). Expression levels of Iba-1, Ikba, IL1B, IL10 and CCL2 significantly increased with time, but independently of FUS intensity. Upregulation of Ikba mRNA levels, a key regulator of NF-kB activation^20^, peaked at 1h post-FUS (∼1.8-fold compared to control), remained elevated at 6h, and returned to near control levels by 24h post-FUS. CCL2, a chemokine implicated in the recruitment of macrophages to damaged tissue^21^ was also upregulated at 1h, peaked at 6h, and remained elevated 24h-post FUS. While a significant effect of FUS-intensity on mRNA levels of the aforementioned pro-inflammatory molecules was not observed, their maximal upregulation occurred at the highest FUS-intensity both 1h and 6h post-FUS (2.6 and 4.48- fold, respectively). In contrast, mRNA levels of the cytokines IL1β and IL10 levels were down- regulated, with their lowest levels (∼0.50-fold of control) detected at 24h and 6h following FUS treatment, respectively.

**Figure 4.**
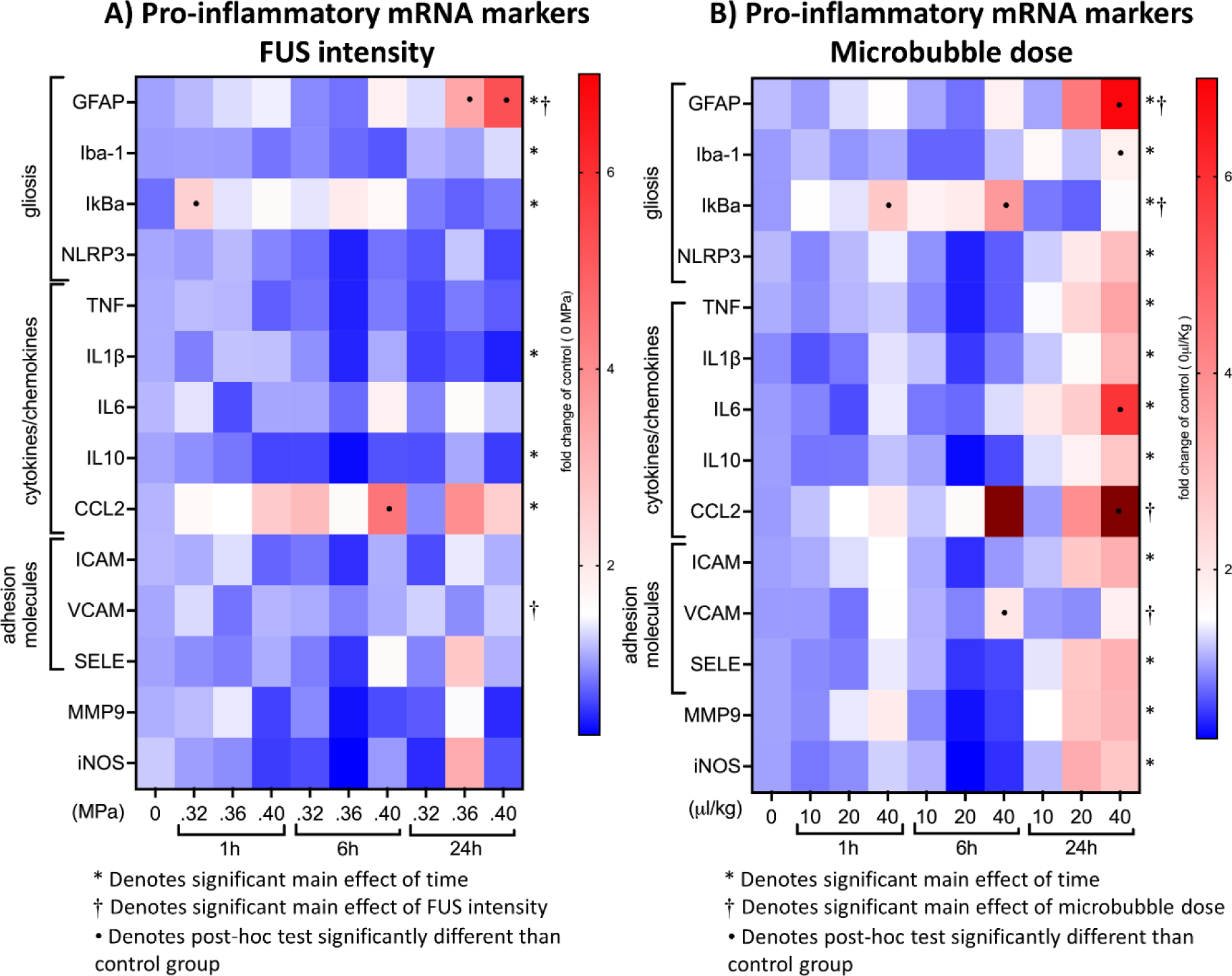
A) Heat map depicting mRNA levels of all pro-inflammatory markers measured by qPCR for the FUS intensity groups. No FUS control is marked as time zero. Color scale shows fold change of control. Based on two- way ANOVA analysis, * denotes a significant effect of the main factor of time, † denotes a significant effect of the main factor of FUS intensity, and • denotes a group significantly different than control following Tukey’s multiple comparison test, all at p<0.05. B) Similar heat map depicting mRNA levels of all pro-inflammatory markers measured by qPCR for the microbubble dose groups.

Overall, greater upregulation of pro-inflammatory mRNA markers was seen with incremental increases of microbubble dose compared to increases in FUS intensity. For almost every mRNA marker measured, there is a significant main effect of increasing mRNA levels with sacrifice time from treatment. GFAP, IKBa, CCL2 and VCAM, also show a significant main effect of microbubble dose, with the highest gene expression at 24h for GFAP (7.2-fold increase) and CCL2 (22-fold) and at 6h for IKBa (3.7-fold) and VCAM (2.0-fold). Iba-1, NLRP3, TNF, IL1B, IL6, IL10, ICAM, MMP9 and iNOS all displayed the highest upregulation for the highest microbubble-dose at 24h post-FUS, albeit the main effect of microbubble dose did not reach statistical significance. Significant differences between sex in the inflammatory response to increased microbubble dose resulted in large data variability, specially at the 24h time point, which may have contributed to the lack of additional statistical significance. An in-depth analysis of sex differences was performed as presented below.

Analysis of correlation was performed separately for FUS intensity and microbubble dose groups for all mRNA against both area of BBB opening and area of microhemorrhage. A Pearson correlation coefficient was calculated for data pooled over time points and FUS intensities/microbubble doses. The area of BBB opening and area of microhemorrhage were strongly correlated for both FUS intensity (r = 0.77, p<0.0001) and microbubble dose (r = 0.66, p<0.0001) groups. All correlations for pro-inflammatory mRNA markers that reached statistical significance are listed in **Table 2** with respective r and p-values. While statistical significance indicates a rejection of the null hypothesis that there is no correlation between the variables, the strength of correlation varies from relatively weak for markers such as IKBa (r = 0.30) and ICAM (r=0.29), to moderate for markers such as VCAM (r=0.54) and strong for CCL2 and hemorrhage (r=0.68).

**Table 2.** Correlation of hemorrhage and BBB opening with mRNA markers of inflammation. Pearson correlation coefficients (r) calculated for FUS intensity groups (data pooled over time points and FUS intensities) and Microbubble dose groups (data pooled over time points and microbubble doses). Only correlations that met statistical significance criteria are listed.

| FUS Intensity Groups |  | Microbubble Dose Groups |  |
| --- | --- | --- | --- |
| BBB Area | Hemorrhage Area | BBB Area | Hemorrhage Area |
| **GFAP**<br>$r = 0.53$<br>$p < 0.0001$ | **GFAP**<br>$r = 0.44$<br>$p = 0.0008$ | **GFAP**<br>$r = 0.60$<br>$p < 0.0001$ | **IKBa**<br>$r = 0.30$<br>$p = 0.026$ |
| **CCL2**<br>$r = 0.57$<br>$p < 0.0001$ | **CCL2**<br>$r = 0.68$<br>$p < 0.0001$ | **CCL2**<br>$r = 0.47$<br>$p = 0.0003$ | |
| | | **TNF**<br>$r = 0.30$<br>$p = 0.028$ | |
| | | **IL6**<br>$r = 0.47$<br>$p = 0.0003$ | |
| | | **ICAM**<br>$r = 0.29$<br>$p = 0.03$ | |
| | | **VCAM**<br>$r = 0.54$<br>$p < 0.0001$ | |

### Sex differences

Analysis of the mRNA data revealed significant differences due to sex in the upregulation of several cytokines and chemokines for treatments done with the 40 µL/kg microbubble dose at FUS-intensity of 0.36 MPa. **Figure 5** shows mRNA levels for those groups with two-way ANOVA analysis done with factors for time and sex. There were no differences due to sex for either the area of BBB opening or the area of microhemorrhage. There were also no differences between male and female mice for two pathognomonic markers of gliosis, GFAP and Iba1. However, female mice exhibited a more robust upregulation at all time points of mRNA levels of the inflammasome marker NLRP3 as well as of all of the cytokines, chemokines, and adhesion molecules that were evaluated, with the exception of VCAM. Many of the differences between females and males were strongly significant, especially at 24 hours. The one marker that showed significantly greater upregulation in male mice compared to female was Ikba.

**Figure 5.**
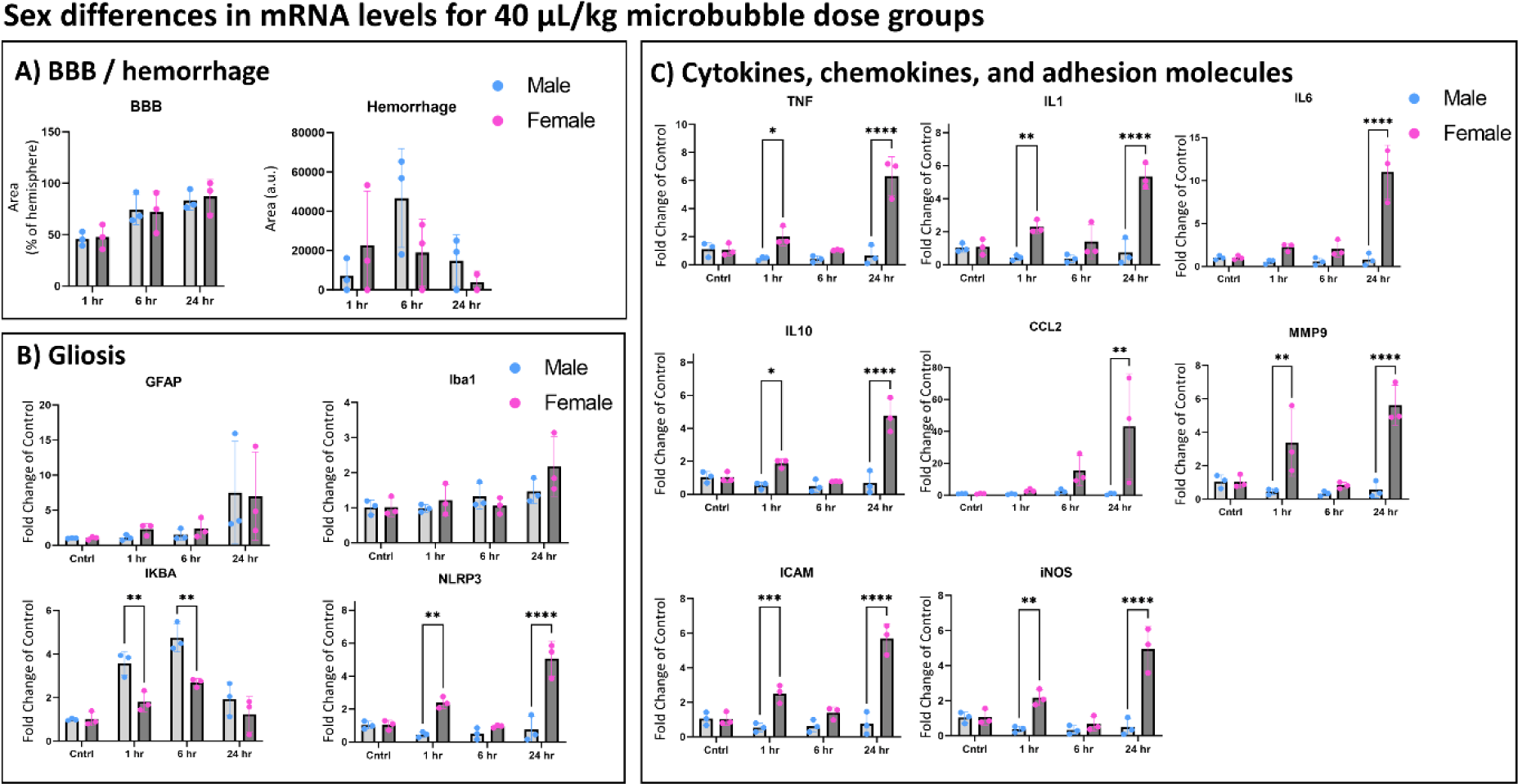
Bar plots show comparisons of mRNA levels in male vs female mice from the 40 µL/kg microbubble dose groups. A) Measures of BBB opening area and microhemorrhage area do not show any differences between the sexes. B) The primary markers for astrocytosis (GFAP) and microgliosis (Iba1) do not show any differences between the sexes. NLRP3 does show significantly greater elevation in female mice compared to male mice, while IKBa is an outlier showing greater elevation in male mice. C) Of the nine cytokine, chemokine and adhesion molecule markers measure, eight showed significantly greater elevation in female mice compared to male mice for at least one time point. Seven of the markers had significance of p < 0.0001.

At the protein level, western blot analysis showed no statistically significant differences in GFAP levels between males and females for the 40 µL/kg microbubble dose groups, which agrees with the qPCR data (**Figure 6A**). Similar to GFAP, ICAM-1 protein expression also increased by 24 hours after FUS and was more robust in females vs. males, albeit remained non-significant (**Figure 6B**). Discrepancy between a significant upregulation of ICAM mRNA but not protein levels could simply relate to the delay between transcription and translation or implicate other post-transcriptional regulatory mechanisms. Expanding the study to include later time points will help resolve this question.

**Figure 6.**
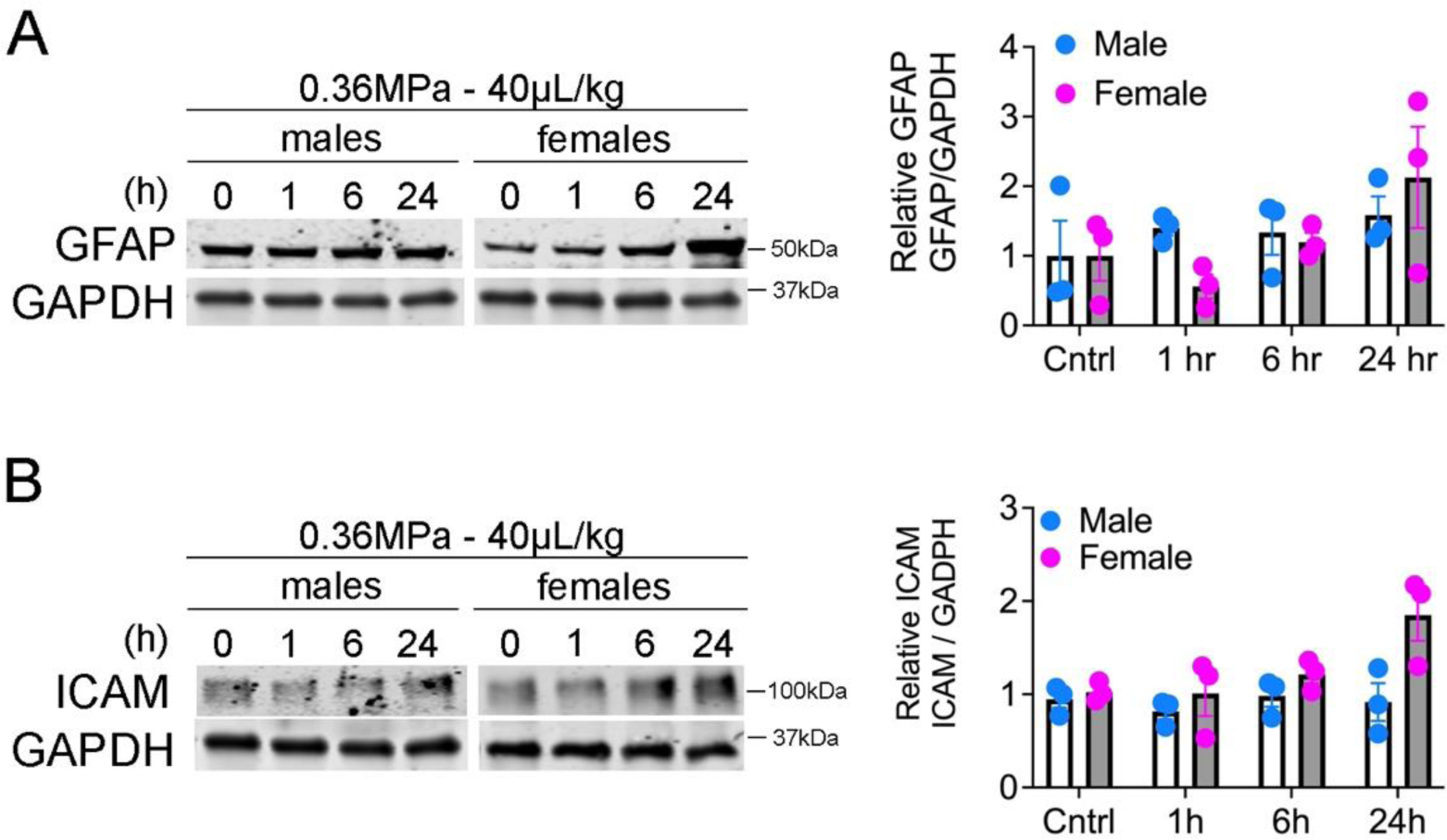
Western blot analysis of GFAP (A), ICAM (B) levels in male vs female mice from the 40 µL/kg microbubble dose groups. Bars represent densitometric quantification expressed as fold change relative to control (n=3 per time point per sex).

### Protective molecules

In addition to pro-inflammatory markers, we also measured mRNA levels of several markers that are involved in tempering down inflammation and restoring homeostasis in endothelial cells (EC). The 3 markers that were selected for this screen include endothelial nitric oxide synthase (eNOS), a key promoter and biomarker of EC health^22–25^, A20, a potent anti-inflammatory and anti-apoptotic protein in EC^26–29^, and VEGF, a key EC growth factor that also promotes EC and neuronal survival^30,31^. **Figures 7A and 7B** show heat maps depicting changes in mRNA levels of these protective markers in response to incremental FUS intensities or microbubble doses. Significant main effects and significant individual group differences based on two-way ANOVA analysis are reported. Most of these markers show significant upregulation with increased FUS intensity and microbubble dose. Albeit, on the individual level, this increase over control was only significant in two instances for A20 and eNOS mRNA levels at 6 hours and 24 hours, respectively, following the 0.36 MPa in the FUS intensity group.

**Figure 7.**
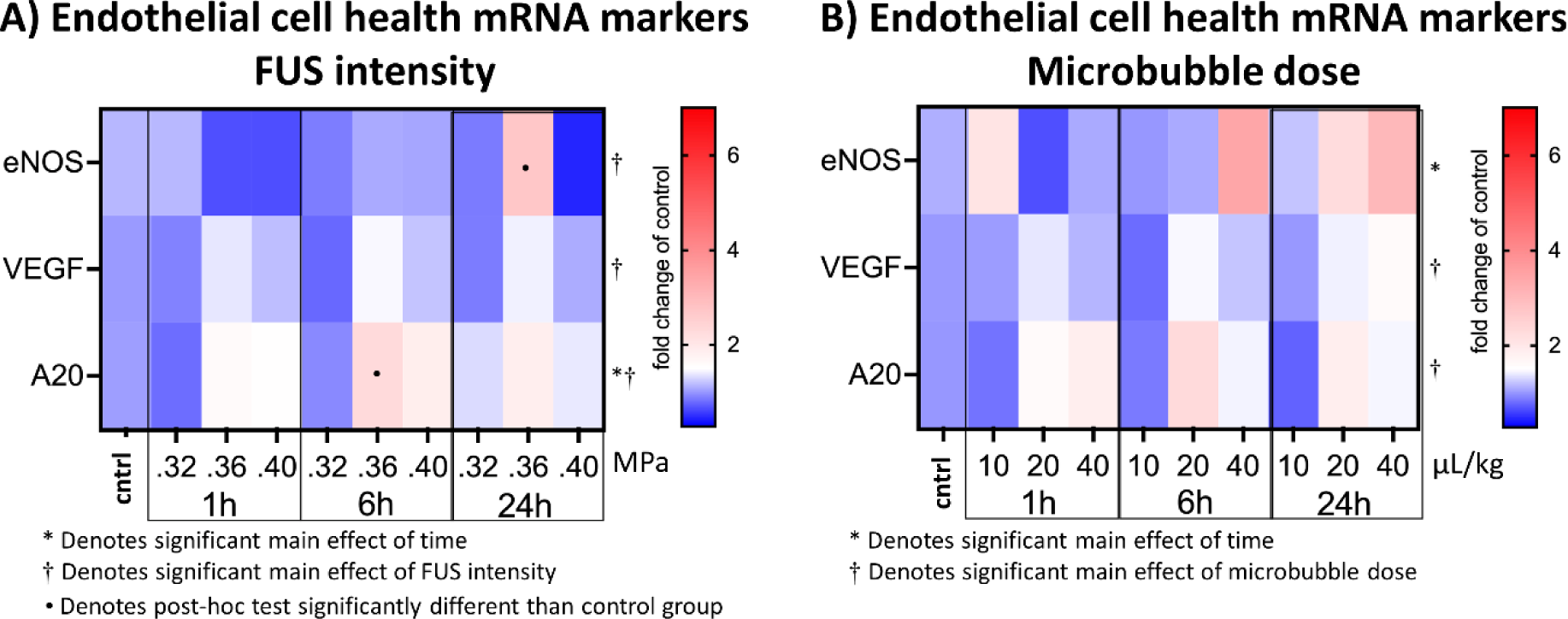
A) Heat map depicting mRNA levels of all anti-inflammatory markers measured by qPCR for the FUS intensity groups. No FUS control is marked as cntrl. Color scale shows fold change of control. As in Figure 4, * denotes a significant effect of the main factor of time, † denotes a significant effect of the main factor of FUS intensity, and • denotes a group significantly different than control following Tukey’s multiple comparison test, all at p<0.05. B) Similar heat map depicting mRNA levels of all anti-inflammatory markers measured by qPCR for the microbubble dose groups.

### Multiple Treatments

To further investigate the pro- and anti-inflammatory responses, an additional group of 8 male mice underwent five FUS treatments, spaced by 1 week, using as FUS parameters a 0.32 MPa FUS intensity and a 20 µL/kg microbubble dose with a sacrifice time 24 hours after the final treatment. **Figure 8** depicts a comparative analysis of mRNA levels of the select panel of pro-inflammatory and anti-inflammatory biomarkers between brains of mice treated 1x vs 5x with FUS. Our results show no differences between groups for either markers of gliosis (GFAP and Iba1) or any of the cytokines or chemokines that were assayed. However, we noted a significantly higher mRNA levels of the protective molecule eNOS following 5x treatments compared to the 1x treatment mice (p< 0.01). This novel result is suggestive of a pre- conditioning effect, which by enhancing EC’s protective armamentarium increases the safety of iterative FUS treatments.

**Figure 8.**
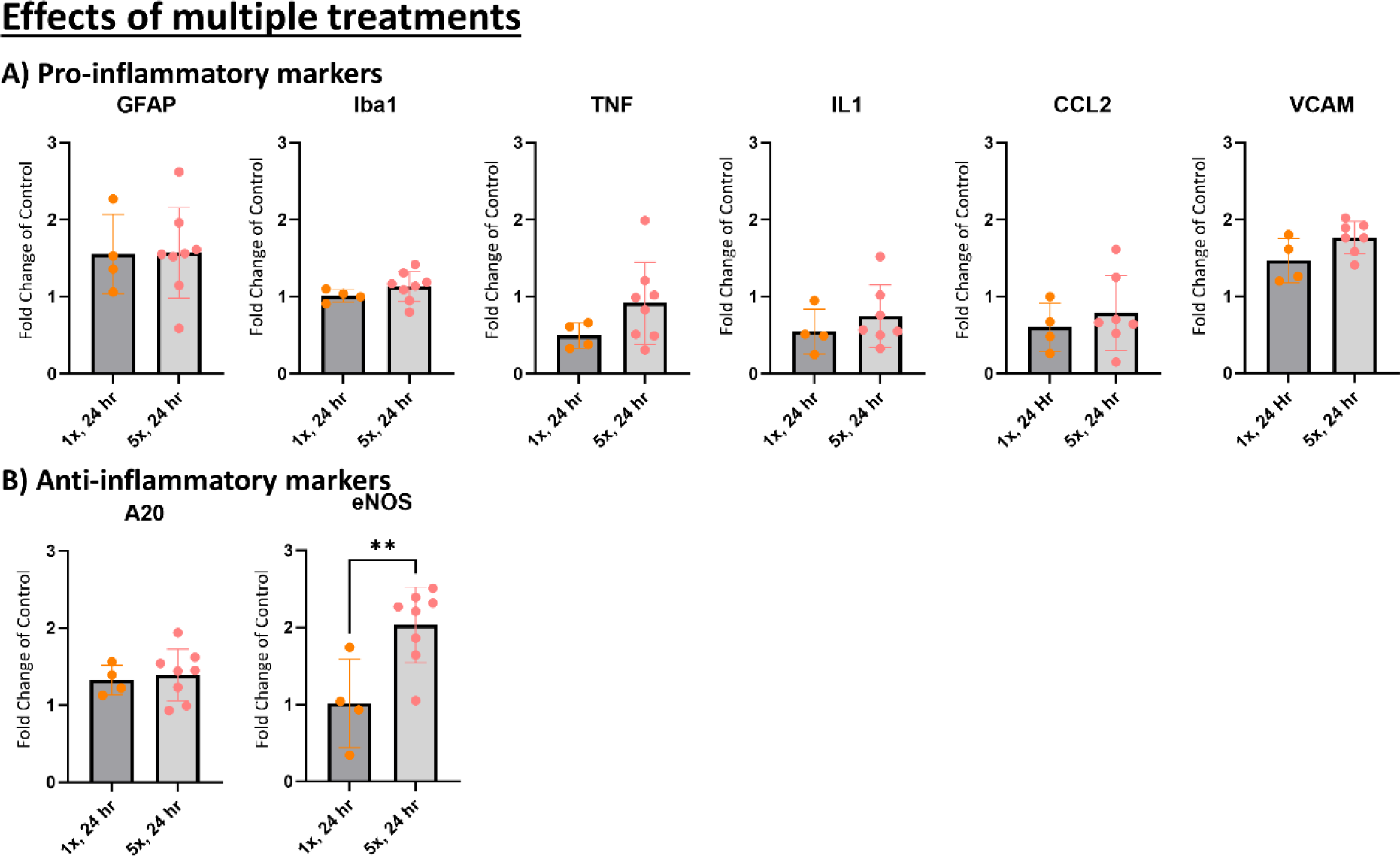
5x vs 1x FUS treatments. A) Select examples comparing mRNA levels of pro-inflammatory markers in male mice after 1x or 5x FUS treatments, with sacrifice at 24 hours after final FUS treatment. Of the 13 pro-inflammatory markers measure, none showed a significant difference between the 1x and 5x groups based on a non-paired two-tailed t-test. B) Similar comparisons for mRNA levels of anti-inflammatory markers. Only eNOS was significantly higher in the 5x treated group compared to the 1x treated group. ** p<0.01

## Discussion

### The inflammatory response as a function of FUS treatment parameters

Results from this study were presented according to temporal progression of the canonical model of brain inflammation: an insult to tissue in the form of BBB disruption; subsequent activation of the brain’s resident glial cells, namely astrocytes and microglia; ensuing production of pro-inflammatory cytokines (*e.g.*, TNFα, IL1, IL6), chemokines (*e.g.*, CCL2) and increased expression of adhesion molecules (*e.g.*, ICAM, VCAM); and ultimately upregulation of anti-inflammatory molecules to restore homeostasis (*e.g.*, eNOS, A20). The initial insult to the brain is FUS-mediated disruption of the BBB, characterized by microbubble oscillation that stretches and constricts blood vessels leading to leakage of blood products into the brain parenchyma, including serum albumin and sometimes red blood cells. The FUS parameters tested resulted in a range of mild to widespread BBB disruption with no to moderate microhemorrhage. We had evidence for activation of the brain’s resident glial cells, namely astrocytosis (GFAP upregulation) at the more aggressive FUS parameters, but only moderate signs of microgliosis (Iba1 upregulation), which is typically considered the main driver of the inflammatory response. While Iba-1 upregulation was considered a marker of microglia activation, recent research suggests that Iba-1 immunostaining is mostly useful to identify changes in microglia morphology and distribution, which is more indicative of activated microglia in the brain than increased expression of Iba-1 per se. Accordingly, our immunostaining using both Iba-1 and TMEM119 (a type I transmembrane protein specifically expressed by resident microglia) showed changes in microglia morphology consistent with activation as early as 1h after FUS, before any increase in Iba1’s expression (Figure S2). Consistent with the observed astrocytocis and microgliosis, we detected an increase in cytokine and chemokine levels following treatments with higher microbubble doses at the 6- and 24- hour time points. Furthermore, there was evidence that an anti-inflammatory response was elicited in reaction to the stronger pro-inflammatory response observed in the high dose microbubble groups, as evidenced by increased expression levels of A20 and eNOS.

These main results agree with the overall picture of inflammation following FUS-BBB opening, as described in the six previously published papers listed in Figure 1. The strongest consensus point being that the inflammatory response occurs when microbubble doses used are well above the clinical dose of 10 µL/kg. All five papers that used a microbubble dose of 100 µL/kg or greater (equivalent Definity dose) reported a significant upregulation of various inflammation markers, even at FUS intensities with a mechanical index lower than that used in this study^10,11,13,15^. Gorick et al. used an equivalent microbubble dose (close to the 40 µL/kg) to the one we used in this study, and reported significant upregulation of GFAP, adaptive immunity markers, and chemokine signaling markers^14^. Interestingly, akin to our study, they also did not observe an increase in Iba1 levels^14^. In addition, there was broad agreement regarding the specific inflammatory markers that are upregulated. Of the four studies that measured levels of specific cytokines, chemokines and adhesion molecules, all saw increases in TNF and IL1, and three out of four noted increases in IL6, CCL2, and ICAM expression levels.

One area of inconsistency among studies relates to the kinetics of the pro-inflammatory response. Five of the studies report significant upregulation of pro-inflammatory markers at the earliest measured time points, ranging from 0.08 to 6 hours post-treatment, with Kovacs et al. indicating that the pro-inflammatory response has largely subsided by 24 hours. However, our data and the results from Ji et al. indicate a mild inflammatory response at 6 hours, escalating to its peak level by 24 hours, at least among the times that were tested. In fact, the Gorrick et al. study also showed inflammation persisting beyond 24 hours.

By assessing a range of FUS intensities and microbubbles doses, we were able to gauge the different effects that result from modifying these treatment parameters. Our results indicate that increases in FUS intensity drive microhemorrhage, while increasing microbubble doses drive inflammation. **Figure 9** illustrates extent of BBB opening, and levels of microhemorrhage and pro-inflammatory markers at the lowest (0.32 MPa / 10 µL/kg, red markers) and highest (0.40 MPa / 40 µL/kg, black markers) settings for each parameter tested. At the lowest parameter settings, BBB opening is evident, yet there is no microhemorrhage and no significant increase in inflammation. When increasing to the strongest FUS intensity, more microhemorrhage is observed compared to the highest microbubble dose group, despite slightly less BBB opening. However, despite worse microhemorrhage, most inflammatory markers remain at baseline levels, and the two that are still upregulated (CCL2 and GFAP) remain at lower levels than what was seen for the high microbubble dose group. Conversely, the high microbubble dose group exhibited slightly enhanced BBB opening, less microhemorrhage, and significantly greater upregulation of inflammatory markers. These results point to the possibility that the inflammatory response in this case might be driven by a form of damage other than the blood products entering the brain parenchyma following BBB opening. For instance, damage signals may be transmitted to EC via mechanosensitive channels triggered by stretching caused by microbubble cavitation.

**Figure 9.**
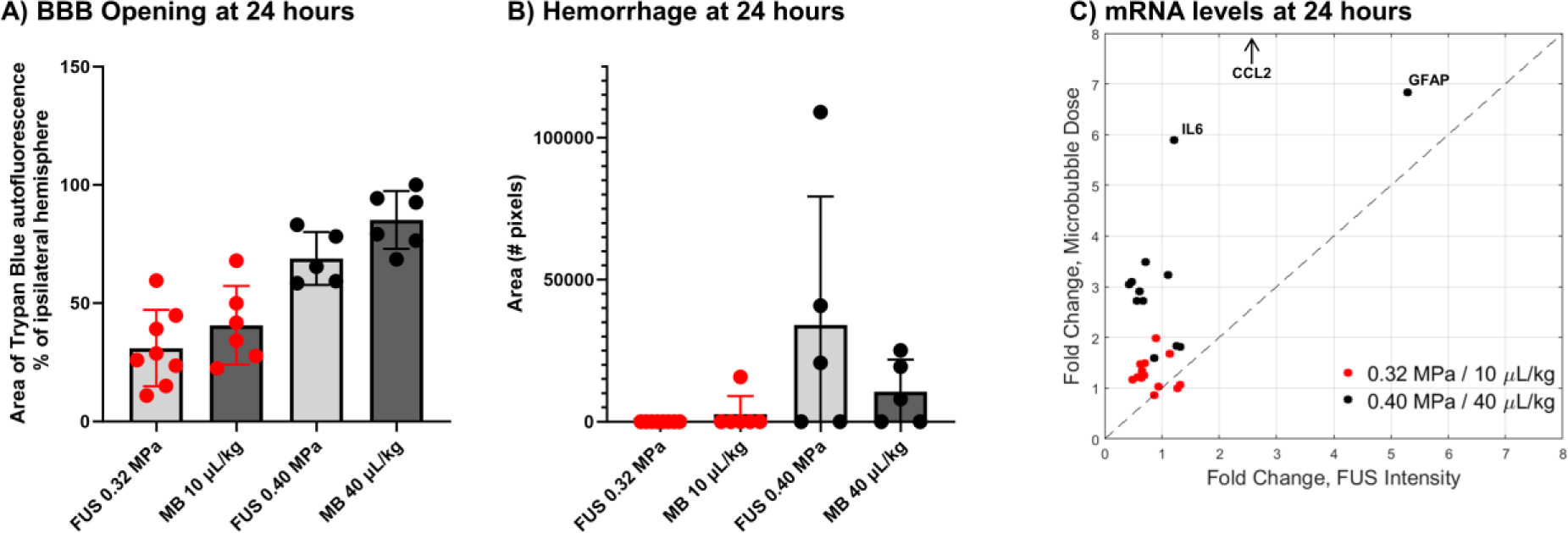
Differential effects of FUS intensity and microbubble dose on BBB opening, microhemorrhage and mRNA levels at 24 hours post treatment. Area of BBB opening (A), area of microhemorrhage (B) and upregulation of pro-inflammatory makers (C) are compared for 24-hour groups treated at the lowest and highest levels of FUS intensity and microbubble dose. The graph in (C) shows each pro-inflammatory marker plotted as fold change for the low/high FUS intensity group on the x-axis and fold change for the low/high microbubble groups on the y-axis.

### Differences due to sex

The literature on sex differences in neuroinflammation is complex and sometimes contradictory. In humans, sex is a known risk factor for various neurological disorders that share neuroinflammation as a central pathophysiological component, a prominent example being Alzheimer’s disease whose prevalence in individuals over 65 years is 2 to 3 times greater in women than men^32^. Sex-related differences in microglia number and morphology have also been observed. However, the mechanistic basis linking these differences to sexual dimorphism in neuroinflammation and disease remain poorly understood.

Pre-clinical studies that attempted to elucidate the mechanisms behind sex-related differences in the context of neuroinflammation often found that microglia activation in response to injury was greater in male mice^33^. For example, two papers that explored neuroinflammatory responses following traumatic brain injury caused by controlled cortical impact injury in wild type mice found that the inflammatory response was more severe in male versus female mice, primarily driven by differences in microglia activation^34,35^.

In our study, we noted a strong and consistent signal indicating that female mice develop greater inflammation in the brain at the highest microbubble dose, as indicated by higher mRNA levels of pro-inflammatory cytokines and chemokines following BBB opening. This difference was evident for almost all cytokine and chemokine markers that were measured and occurred in all groups and at the three time-points tested, making it unlikely that it was an artifact of treatment day or tissue batch analysis. Interestingly, the degree of BBB opening, extent of microhemorrhage and severity of astrocytosis (GFAP) and microgliosis (Iba1) were comparable between males and females, suggesting that the molecular mechanism(s) responsible for male-female differences lie downstream of these factors. Additional experiments are needed to clarify the molecular basis of sex-related differences, including hormonal influences.

### Role of protective molecules

In this study we also investigated three molecules that are involved in the regulatory response to injury and inflammation, A20, eNOS and VEGF. These molecules primarily function to contain inflammatory responses, prevent apoptosis, support reparative and regenerative mechanisms, and restore homeostasis. A20 is a ubiquitous and potent negative feedback regulator of NFκB signaling, suppressing downstream upregulation of pro-inflammatory molecules in the three cellular subtypes of the neurovascular unit affected by FUS/MB disruption of the BBB: endothelial cells, astrocytes/microglia and neurons^16,26,27,29,36,37^. A20 also exerts broad cytoprotective and anti-apoptotic effects, preserving survival, particularly in endothelial and neuronal cells survival, in part through inhibiting proteolytic cleavage of caspases 8 and 3 and increasing eNOS expression and activation in EC^22,27,28,38–40^. The dual anti-inflammatory and anti-apoptotic functions of A20 in endothelial and neuronal cells could be highly advantageous in minimizing side-effect of FUS-MB, particularly if higher FUS intensity or microbubble doses are required for clinical benefits^36,41^. Endothelial NOS is an endothelial-cell specific enzyme that regulates endothelial nitric oxide (NO) production, maintaining vascular integrity and tone. VEGF, mostly the VEGF-A isoform, primarily produced by endothelial and glial cells in response to hypoxic and pro-inflammatory signals, promote endothelial and neuronal cell growth and survival^23,24,42–47^. Both eNOS and VEGF, through nitric oxide/cGMP dependent mechanisms, increase endothelial permeability, including at the BBB level^48–51^. Controlled and transient increase in permeability could synergize with FUS/MB treatment, whose mechanisms of BBB disruption relate to targeting tight junction components such as occludins, claudins, and Zona Occludens-1 (ZO-1and ZO- 2), enhancing efficacy.

Our results reveal time-, FUS intensity- and microbubble dose-dependent modulation of A20, eNOS and VEGF mRNA levels in mouse brains following FUS/MB treatment. The expression of these molecules mostly paralleled that of the pro-inflammatory markers, underscoring their role in balancing procedure safety. While changes in VEGF mRNA levels were mild, significant upregulation of A20 and eNOS was observed in response to FUS. A20/TNFAIP3 and eNOS mRNA levels were maximally and significantly elevated above baseline controls at 6 and 24 hours, respectively, in the 0.36 MPa FUS intensity group. However, their levels notably decreased when the FUS intensity was increased to 0.4 MPa. Specifically, A20 mRNA levels at 6 hours in the 0.4 MPa group only showed a moderate increase over controls, whereas the 24-hour eNOS mRNA levels were actually lower than control levels, indicating endothelial cell dysfunction and potential toxicity. These results suggest that tolerance to higher FUS intensity may be compromised by inadequate protective responses, highlighting the need for establishing safety thresholds. Additional experiments testing FUS intensities between 0.36MPa and 0.4MPa are planned to identify the cutoff.

In contrast to the curve drop-off observed at the highest FUS intensity, mRNA levels of the A20/eNOS protective module increased in a microbubble dose-dependent manner up to the highest dose tested. This suggests a broader therapeutic window for microbubble dose, and implies that increased microbubble dosing to possibly enhance BBB disruption and delivery of therapeutics might be safer for future clinical applications. Additional experiments are required to identify the maximal microbubble dose that could be safely applied without toxicity.

### The effect of multiple treatments

To assess the impact of multiple FUS-BBB opening treatments, we selected a conservative parameter set of 0.32 MPa FUS intensity and 20 µL/kg microbubble dose, which caused BBB opening without eliciting an inflammatory response after 1x treatment. We first determined whether these conservative FUS parameters applied in relatively close succession, with only a 7-day interval between treatments, would provoke an inflammatory reaction. Analysis of pro-inflammatory markers indicated that this was not the case. Indeed, there was no significant increase in pro-inflammatory mRNA levels above the 1x treated mice, with the exception of VCAM whose mRNA levels showed a non-significant trend for increase (p = 0.079). Next, we tested the hypothesis that repeated low-level BBB opening could “pre-condition” the brain microenvironment by promoting the upregulation of the protective aspects of the response to inflammation and injury. The concept of ischemic or inflammatory preconditioning to promote tissue resistance to injury has been implemented in other fields such as cardiac, lung, liver and kidney surgeries, transplantation, and even in the context of inflammatory brain diseases^22–24^. Remarkably, eNOS mRNA levels in brains of 5x treated mice were significantly higher compared to the 1x treated mice. However, we did not see differences in A20 levels, at least at the time points that were examined.

Altogether, these results suggest that the pro-inflammatory response to FUS-BBB opening is mostly commensurate to the intensity of the insult, but not necessarily activated by multiple sub-threshold events, with the exception of eNOS. We surmise that this pre-conditioning regimen, which increases basal levels of eNOS, may enhance EC resistance to injury, potentially improving the safety of higher FUS intensities that may be necessary for therapeutic benefits. Alternatively, increased eNOS expression could also augment NO availability, synergizing with FUS/MB to boost BBB permeability and improve efficacy and delivery of therapeutics, all while implementing lower/safer FUS intensity levels and microbubble doses. This warrants additional investigation. Future studies are aimed at expanding our kinetics analysis after FUS BBB opening to better assess the granularity of the general and anti-inflammatory protective response to single vs. multiple treatments, both at the transcriptomics and proteomics levels.

Notably, an original version of the multi-treatment study was performed with Optison microbubbles at a dose of 100 µL/kg, which is twice the clinical dose and roughly equivalent to 20 µL/kg of Definity. Unfortunately, several of the healthy wild type mice died after the second and third FUS-BBB opening treatments, likely because of an anaphylactic response to the human albumin contained in Optison.

## Acknowledgements

The research was supported by NIH grants R21 EB030173 and R01 NS123557.

