## Supplementary Figures for "Characterization of focused ultrasound blood-brain barrier disruption effect on inflammation as a function of treatment parameters"

### Supplementary Material

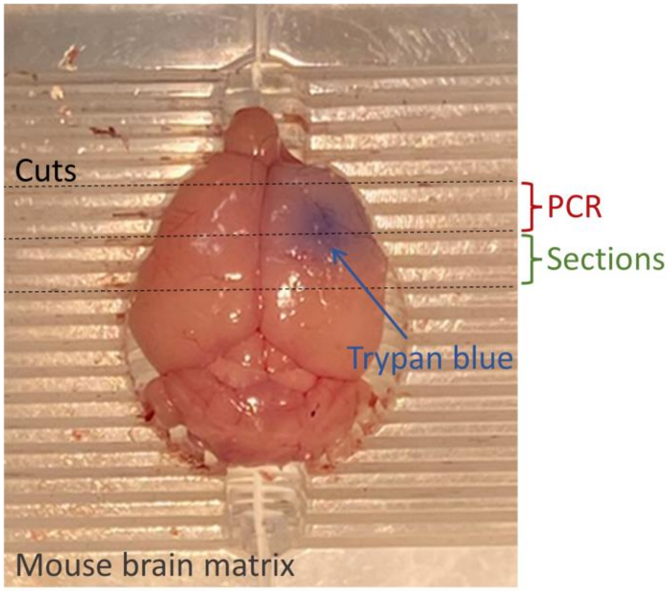


***Figure S1***. *Fresh mouse brains were dissected in a mouse brain matrix to harvest tissue for PCR/WB analysis and sections for IHC. The initial cut was made through the center of BBB opening based on trypan blue signal. An anterior 2mm slab was harvested for PCR/WB and a posterior 2mm slab was harvested to be used for IHC sections.*

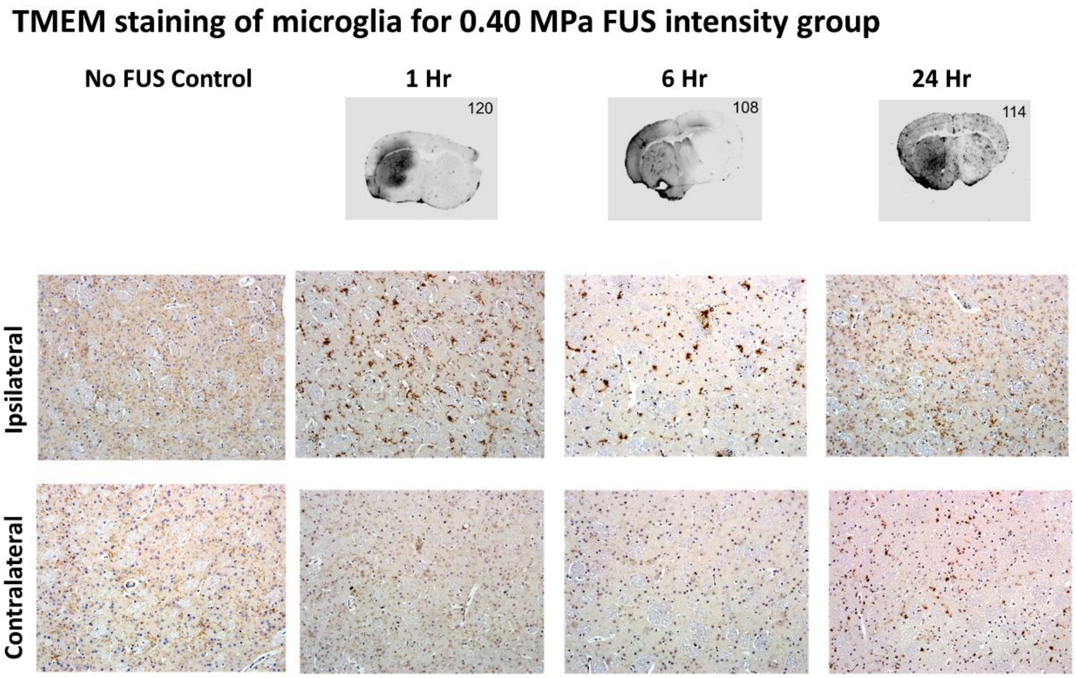


*Figure S2. Example images of TMEM staining for microglia morphology in FUS-targeted (ipsilateral) and non-targeted (contralateral) striatum. Examples are all female mice from the 0.40 MPa FUS intensity group at 1- 6- and 24-hr time points.*
